# EdgeHOG: fine-grained ancestral gene order inference at tree-of-life scale

**DOI:** 10.1101/2024.08.28.610045

**Authors:** Charles Bernard, Yannis Nevers, Naga Bhushana Rao Karampudi, Kimberly J. Gilbert, Clément Train, Alex Warwick Vesztrocy, Natasha Glover, Adrian Altenhoff, Christophe Dessimoz

## Abstract

Ancestral genomes are essential for studying the diversification of life from the last universal common ancestor to modern organisms. Methods have been proposed to infer ancestral gene order, but they lack scalability, limiting the depth to which gene neighborhood evolution can be traced back. We introduce edgeHOG, a tool designed for accurate ancestral gene order inference with linear time complexity. Validated on various benchmarks, edgeHOG was applied to the entire OMA orthology database, encompassing 2,845 extant genomes across all domains of life. This represents the first tree-of-life scale inference, resulting in 1,133 ancestral genomes. In particular, we reconstructed ancestral contigs for the last common ancestor of eukaryotes, dating back around 1.8 billion years, and observed significant functional association among neighboring genes. The method also dates gene adjacencies, revealing conserved histone clusters and rapid sex chromosome rearrangements, enabling computational inference of these features.

## Introduction

Modeling the ancestral genomes corresponding to internal nodes of a species phylogenetic tree is a powerful tool to gain understanding on the chain of genetic events through which genomes diversified. Commonly, this is achieved through ancestral gene repertoire reconstructions, which essentially yield lists of ancestral genes as proxies for ancestral genomes^1,2^. However, these methods do not integrate the information of gene contiguity across genomes and thus cannot inform on events of genomic rearrangements, or lack thereof. Ancestral gene order inferences have then emerged to fill this gap, whether to flag rearrangements associated with speciation or to highlight novel candidate systems of functionally-associated genes through the identification of conserved genomic neighborhoods^3–8^. An impressive milestone was set by Muffato et al.^*3*^ with their *Ancestral Gene Order Reconstruction Algorithm* (AGORA) method and the gene order reconstructions from 98 extant genomes (93 Vertebrates and 5 outgroups), characterizing rearrangement rates across key vertebrate lineages in the process. Yet, state-of-the-art ancestral gene order inference methods such as AGORA typically scale at least quadratically with the number of input genomes and are therefore unable to process large phylogenies encompassing more than hundreds of extant genomes^9^. This limitation affects both the accuracy and evolutionary scope of potential analyses, as including more extant genomes allows for a higher resolution in reconstructing ancestral gene order and tracing their evolutionary histories further back in time.

Here, we introduce edgeHOG, a method to reconstruct ancestral gene orders across large phylogenies while maintaining, and even exceeding, the levels of resolution and predictive accuracy set by AGORA. To achieve this, edgeHOG leverages the framework of hierarchical orthologous groups (HOGs) computable on arbitrary datasets using orthology inference pipelines^10–12^ and made available by orthology databases such as OMA^13^, Hieranoid^14^ or EggNOG^15^ to anchor comparisons of gene adjacencies between genomes and propagate gene order predictions along the species tree. We applied edgeHOG to all 2,845 extant genomes from the OMA database^13^, resulting in 1,133 fully browsable and downloadable ancestral genomes encompassing all three domains of Life. The reconstruction revealed a significant association between gene order and function in the last eukaryotic common ancestor (LECA). The ability of dating gene adjacencies also uncovered intriguing patterns of chromosomal evolution, such as the conserved histone gene clusters in metazoans and the younger gene adjacencies on sex chromosomes across various species. EdgeHOG is available as an open source standalone tool to process arbitrary datasets. In combination with the recently released FastOMA^11^ software, edgeHOG makes it possible to reconstruct ancestral gene orders for thousands of genomes within days.

## Results

### Algorithm overview

EdgeHOG is available as an open source command-line tool distributed as a python package on PyPI and GitHub (https://github.com/DessimozLab/edgehog). It requires a rooted species tree, the coordinates of the genes on the contigs of the species genomes (as GFF files) and the HOGs of the genomes either available for download on various orthology databases in OrthoXML format or computable from proteomes using software such as OMA Standalone package^10^ or FastOMA^11^.

#### Ancestral gene content reconstruction

Conceptually, HOGs can be thought of as ancestral genes, as they encompass orthologs and paralogs descending from a common ancestral gene at a specific taxonomic level (*i.e*. internal node of the species tree)^16–18^. HOGs are nested within HOGs of a higher taxonomic level, thereby modeling the lineage of genes, assuming strict vertical inheritance along the species tree. When distinct HOGs defined at the same taxonomic level are nested in a higher level HOG, they can be thought of as ancestral in-paralogs. EdgeHOG relies on the pyHAM^19^ python package to parse the OrthoXML file of HOGs and map ancestral genes to internal nodes of a phylogenetic tree, gradually leading to gene lineage and ancestral gene content reconstructions (**Figure 1**).

**Figure 1.**
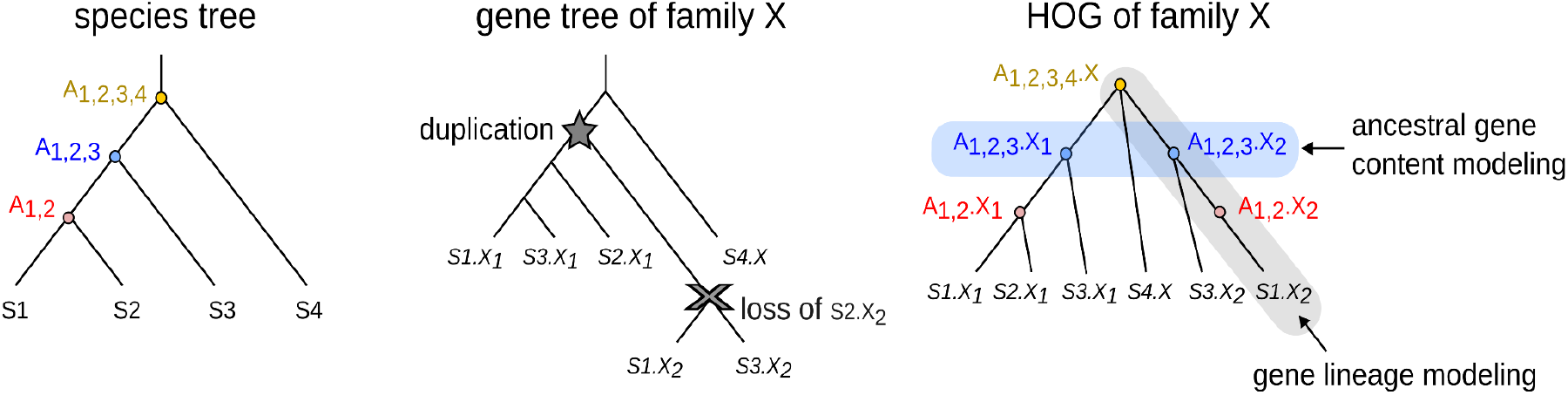
HOGs explicitly model gene lineage, thus ancestral gene contents. The species tree on the left displays the evolutionary relationships between four extant species (S1, S2, S3, S4). Its internal nodes are proxies for ancestors (A_1,2_, A_1,2,3_, A_1,2,3,4_). The central gene tree illustrates an example of inferred evolutionary history for a gene family (coined X) present in all species. The HOG object of the family X can be computed from the reconciled gene tree of X as in EggNOG, or using fast graph-based methods as in OMA. Once pyHAM processes the HOG of family X, the graph on the right is created: each node corresponds to an extant or ancestral gene and each edge links a descendant gene to its parent gene in its most direct ancestor. When paralogs are connected to the same parental gene, it models an event of gene duplication (happening here between ancestor A_1,2,3,4_ and descendant A_1,2,3_). The collection of gene nodes of all HOGs’ graphs at a given ancestral level eventually serves as proxies for its ancestral gene content.

#### Bottom-up propagation of gene adjacencies

Using descendant-to-parent gene links encapsulated in gene lineages, any adjacency observed/predicted between two genes at a given level of the phylogeny is mapped, when possible, to the corresponding parental genes in the upper taxonomic level. When a gene has no parent but its two flanking neighbors have one, an edge between these two neighbors is created and propagated to the upper taxonomic level, thereby modeling events of gene emergence and insertion between two older genes. Eventually, a network is created at each level of the phylogeny, where a node is an ancestral gene and an edge links two genes inferred to be of closest proximity, with a weight equal to the number of propagations from descendant extant genomes (**Figure 2a**).

**Figure 2:**
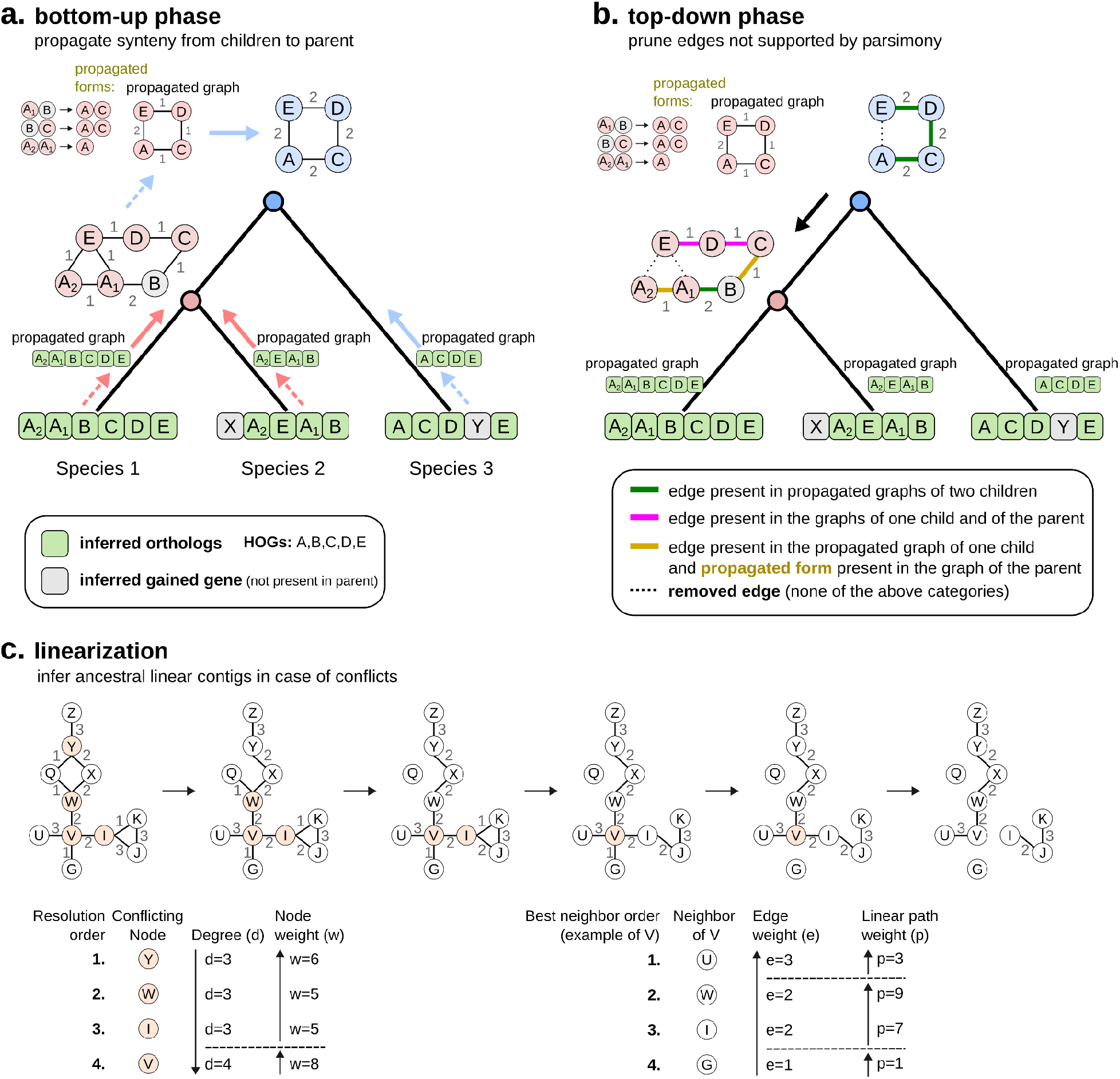
EdgeHOG’s algorithm. **a. Bottom-up phase.** Traversing the guide species tree from leaves to root, an adjacency between two genes is propagated to a direct ancestor as long as it is inferred to have the two ancestral genes. Inferred gene gain (in grey) is accounted for by propagating an adjacency to the parental level only between the two flanking neighbors. All edges propagated to the parental level are summarized in a “propagated graph” and the propagated form of each real edge is stored. **b. Top-down phase**. Traversing the tree from root to leaves, any adjacency not supported by parsimony is removed, that is essentially any edge supported by only one child and not by the parent (hence wrongly propagated before the last common ancestor in which the edge emerged). **c. Linearization phase**. The linearization step flags conflicting genes (degree, i.e. number of edges per node, > 2) and removes edges until the degree of a conflicting gene is no more than 2. The order in which conflicting nodes are resolved and the hierarchy of neighbors for a conflicting node are explained in the bottom of the panel. For each ancestor, the linearized graph constitutes edgeHOG’s main output.

#### Top-down removal of edges not explained by parsimony

A drawback of the bottom-up phase is that when a novel adjacency between two old genes arises through genomic rearrangement, propagating the adjacency to the ancestral level is essentially a mistake. The top-down phase is meant to remove any edge propagated in ancestral synteny networks that is not explained by a parsimonious rationale, namely propagated before the last common ancestor (LCA) in which the adjacency is inferred to have emerged (details in **Figure 2b**). Because the criterion of edge removal does not consider edge weights, the top-down phase is not affected by any potential tree imbalance.

#### Linearization of synteny networks

After these edge removals, some ancestral genes may still have more than two neighbors, due to orthology/paralogy misinferences, incorrect species tree topologies, convergent or reticulate evolution of gene adjacencies. Hence, the linearization step “resolves” conflicting genes by selecting their most likely flanking reconstructible genes based on edge weight (details in **Figure 2c**). This results in linear ancestral contigs at each level of the phylogeny, which altogether compose a so-called ancestral genome.

### Validation on simulated genomes

To evaluate the method in a rigorously controlled experiment, we simulated genome evolution resulting in a clade of 100 short extant genomes of ∼400 genes with gene losses, gene duplications, translocations and inversions using ALF^20^. We then inferred the gene orders of the 99 ancestral levels of the input guide species tree with edgeHOG and AGORA under default parameters, using HOGs computed from the proteomes of the 100 extant genomes (converted into reconciled gene trees for AGORA). For both tools, we compared the predicted gene order at each internal level of the tree with that of the known order from the simulation (**Figure 3a**). Overall, the harmonic mean precision (percentage of predicted adjacencies being correct) and recall (percentage of real adjacencies being predicted) of edgeHOG reached 98.9% and 96.8%, respectively, while that of AGORA reached 96.0% and 94.9%. In addition, edgeHOG’s precision and recall was stable at every depth of the phylogeny, while AGORA’s tended to be lower for more recent ancestors (**Figure 3a**). This somewhat counterintuitive behavior was already reported by the authors of AGORA^3^.

**Figure 3.**
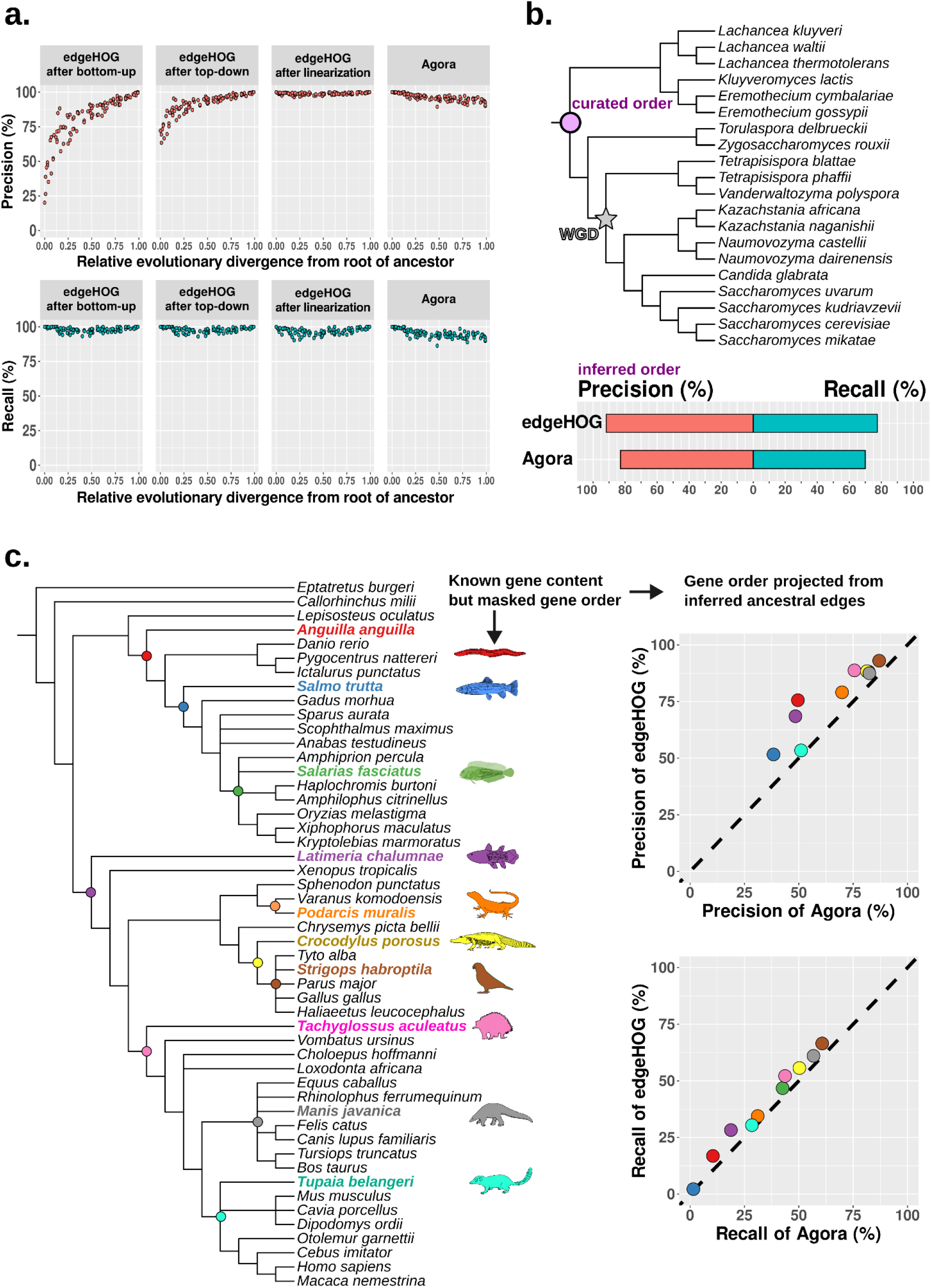
Benchmarks. ***A. Simulated genome evolution benchmark*.** Each dot in each plot corresponds to one of the 99 ancestral levels of the species tree comprising 100 extant genomes. The x-axis gives the *Relative Evolutionary Divergence*^*21*^, which goes from 0 for the root to 1 for the farthest leaf. The y-axis of the top row gives the precision of each algorithmic step of edgeHOG and of Agora, namely the proportion of predicted edges at the ancestral level that are true edges in the corresponding “real” simulated ancestral genome. The y-axis of the bottom row gives the recall, namely the proportions of true edges that are predicted by each method. **b. Yeast Gene Order Browser (YGOB) benchmark**. The species tree on the top corresponds to the phylogeny of the 20 yeast genomes present in the YGOB. The star indicates an event of whole genome duplication that happened in the last common ancestor of a clade of 12 yeasts. The pink circle indicates the root of the tree, for which YGOB-curated ancestral gene order reconstruction is available. This manually curated gene order is compared with edgeHOG’s and AGORA’s predictions, enabling assessment of the precision and recall of both tools. **c. Masked extant gene orders benchmark**. The species tree on the left corresponds to the phylogeny of 50 extant genomes present in the OMA database, sampled to be representative of the diversity of the Vertebrata clade. The 10 colored extant genomes correspond to those whose gene order is masked (and is to be inferred). The 10 colored internal levels correspond to the most direct ancestor of each masked species. The scatterplots on the right compare the performance of edgeHOG with that of AGORA at inferring masked edges using a projection of each edge between two ancestral genes at the parental level onto their corresponding descendant genes in the extant masked species. Note the recall of ∼0% for *Salmo trutta* is because a whole gene duplication happened on the terminal branch, and thus there is no information available to “phase” the duplicated genes.

To gain insights in the relative contribution of edgeHOG’s three algorithmic phases (bottom-up, top-down, and linearization), we also measured precision and recall after each phase; while recall is already high after the first phase, the analysis makes it clear that the top-down and linearization phases are also needed to achieve the high precision, particularly for deep ancestors (**Figure 3a**).

### Validation on real data: Yeast Gene Order Browser benchmark

To assess edgeHOG on real data, we took advantage of the expert and thorough work performed by the Yeast Gene Order Browser (YGOB) to manually curate the likely gene order in the last common ancestor of a clade of 20 yeast species^22^. We ran edgeHOG and AGORA on the dataset and compared the predicted adjacencies at the root of the species tree with that annotated by the YGOB. Despite the complex evolutionary history of the 20 species, involving an ancient hybridization between two ancestors, followed by whole genome duplication in the last common ancestor of a clade of 12 yeasts^23^, the precision and recall of edgeHOG reached 91.7% and 77.4%, respectively, while that of AGORA reached 82.8% and 70.0% (**Figure 3b**).

### Validation on real data: inferring masked extant gene orders from ancestral gene order predictions

Despite the well-established nature of YGOB and its curated ancestral gene order, its focus on a specific yeast clade and evolutionary divergence raises questions about the generalizability of our findings. Additionally, there is a possibility that YGOB might favor the method that has the most in common with its inference process. We thus designed another test on real data encompassing a diverse set of 50 vertebrate genomes and masked the gene order in 10 genomes, *i.e*. treating each gene as if it were on its own contig, effectively removing any information about the actual order of the genes in these genomes. We ran edgeHOG and AGORA on this dataset to assess their ability to accurately reconstruct the gene order in these 10 genomes using information from the other 40 genomes. Specifically, we inferred the gene adjacencies of each masked genome by mapping the predicted adjacencies of their most direct ancestor onto the corresponding descendant genes in the masked genome. Under the assumption that an ancestral genome shares a substantial number of gene adjacencies with a direct descendant, all else being equal, the number of accurate predictions in a masked genome projected from ancestral edges can be considered as a proxy of the quality of the ancestral gene order inference. The 10 genomes were selected to obtain a diverse set of characteristics, in terms of their taxonomic distribution, proteome quality as assessed by OMArk^24^, number of sister taxa with respect to the direct ancestor, and degree of polytomy of their direct ancestor (**Figure 4, Figure S1a**).

**Figure 4.**
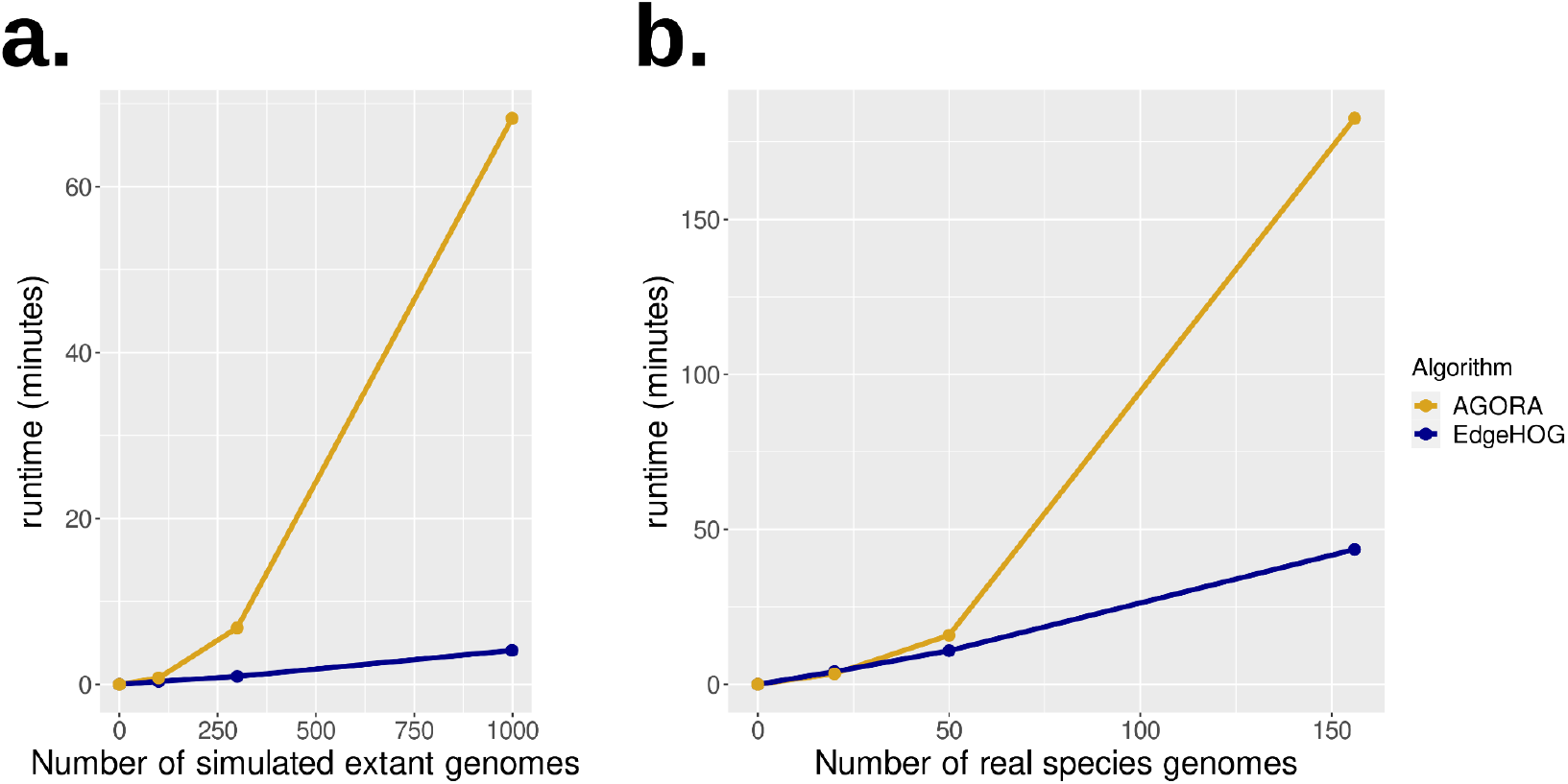
Runtime on one processor as a function of the size of the input phylogeny. **a. Simulated genomes** (mean = 412 genes, standard deviation = 22). **b) real genomes from the Vertebrata clade** (mean = 20,575 genes, standard deviation = 5239).

Once again, for each of the 10 masked species, the performance of edgeHOG met and even surpassed the levels of precision and recall established by AGORA, (**Figure 3c**). Similar results were obtained when processing the full Vertebrata clade of 156 species, keeping the same 10 masked species (**Figure S1b)**.

### Scalability

To empirically assess computational efficiency and scalability, we measured runtime as a function of the size of the input phylogeny (**Figure 4**). EdgeHOG’s runtime scales almost linearly with the size of the input dataset, a performance achieved through the post-order and pre-order traversal of the species phylogeny to create and remove edges in synteny networks, as opposed to pairwise comparisons of extant gene orders, which increases quadratically. In the current implementation, the slight increase in the slope of edgeHOG is almost entirely due to writing the output—ancestral genes being currently named as a concatenation of all descendant genes (which becomes bigger as the size of the phylogeny increases).

### Large collection of ancestral genomes across the three domains of Life in the OMA browser

The scalability of edgeHOG enabled us to process the entire OMA orthology database, currently spanning 2845 extant genomes (1965 bacteria, 173 archaea, 707 eukaryotes), in less than 4 hours on a single processor. To our knowledge, this represents the first attempt to infer ancestral gene orders at the tree of life scale. The resulting collection of 1133 ancestral genomes represents a unique resource to study ancestral synteny across clades of all three domains of Life. The interface to browse these data is described in the latest OMA paper^13^.

In Figure 5, we show an example of a reconstructed ancient ancestral genome with the gene order inference for the last eukaryotic common ancestor (LECA). Specifically, edgeHOG inferred 1009 ancestral contigs in LECA (**Figure 5a**). The functional descriptions of HOGs (ancestral genes) from the same contigs gives credit to the inference since neighboring genes often appear functionally related, in line with the notion that there is a link between genetic linkage and functional association^25^ (**Table S1, Materials and Methods**). The GO enrichment analysis of contigs (GO terms of genes of a contig as foreground, GO terms of genes of the ancestral genome as background) confirms this trend by highlighting 194 contigs enriched in ancestral genes contributing to the same biological process (Fisher’s exact test, Bonferroni-corrected p-value < 0.05). As a sanity check, we repeated the analysis after randomizing gene order (preserving contig size and GO annotations) and the number of functionally enriched contigs was indeed much lower (mean=14.6, stdev=7.9) (**Figure 5b**, detailed procedure in **Materials and Methods**). Remarkably, the reconstructed contigs contain genes that altogether summarize core pathways, with primary metabolism, translation, DNA repair, and stress responses being the most represented categories of pathways (**Figure 5a, Table S1**). However, a few contigs are erroneous based on our knowledge of eukaryotic evolution, such as contigs containing genes involved in photosynthesis, likely induced by the cyanobacterial ancestry of chloroplastic gene adjacencies (**Discussion**). This highlights the potential for future algorithmic improvements, notably accounting for the possibility of reticulated evolution. Finally, we estimated that extant eukaryotic genomes possess between 0 to 4.6% (1.25% on average) of the gene adjacencies inferred in LECA (**Figure 3c, Table S2**), the most conserved adjacency being between Histone 2A and Histone 2B (in 66% of all tested extant eukaryotes) (**Table S2**).

**Figure 5.**
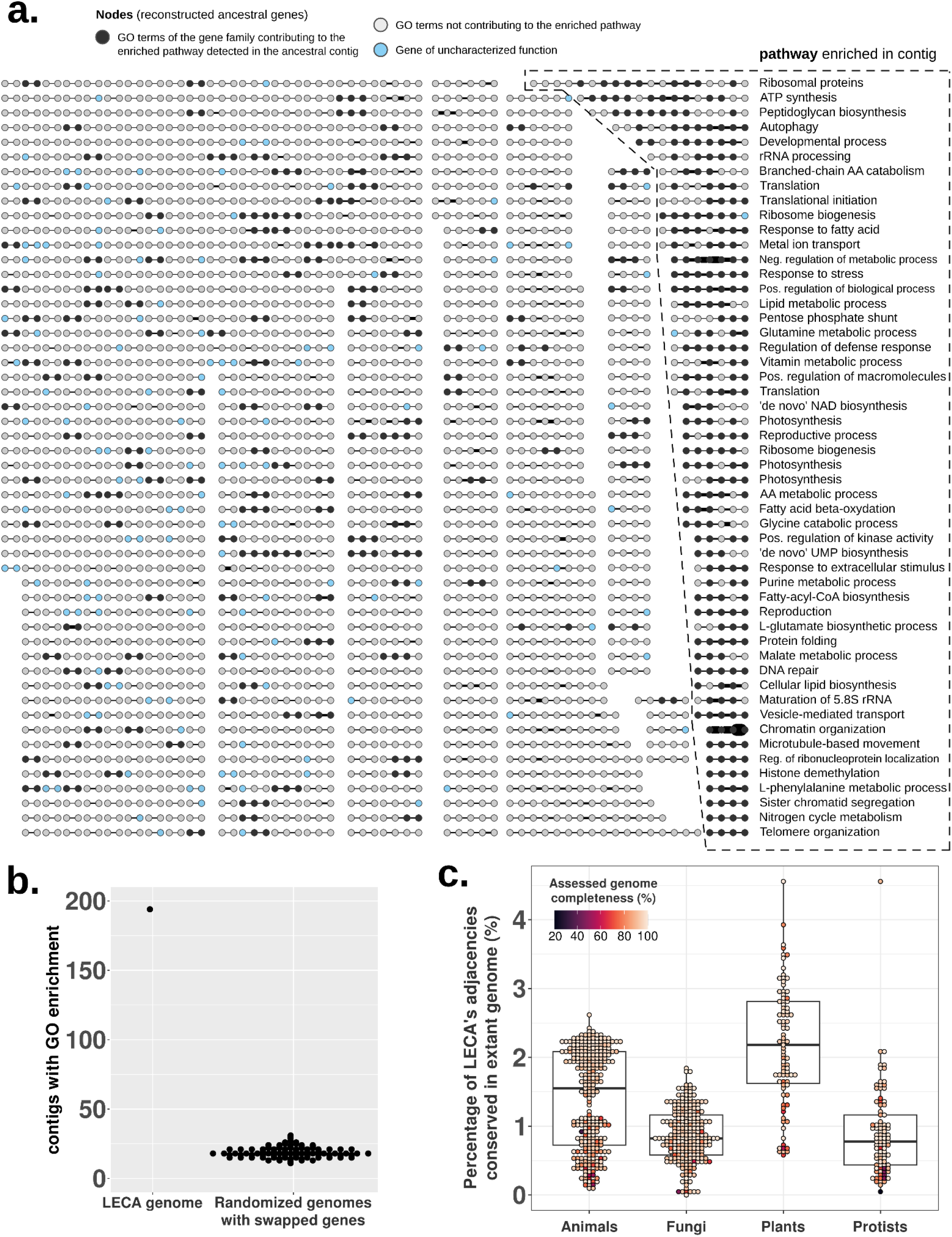
Functional analysis of the 1009 ancestral contiguous regions inferred by edgeHOG in the last eukaryotic common ancestor (LECA). **a) Landscape of ancestral contigs.** Each dot represents an ancestral gene present in a reconstructed contiguous region. Each edge links two genes inferred to be mutually closest to each other among the pool of reconstructible genes. The thickness of an edge is proportional to the square root of its weight, ranging from 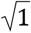 to 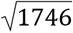. The weight of an edge corresponds to the number of times the edge was propagated from descendant extant genomes, thus depicting the level of conservation of genomic neighborhoods. Within each contig, a black dot corresponds to a gene whose descendant genes’ GO terms contribute to the enriched pathway detected in the contig. A contig with only black dots implies therefore that all of its genes have been detected as participating in the same biological pathway. A blue dot means that the function of the gene family is uncharacterized. For the sake of readability, enriched pathways are only written for contigs displayed at the right border of the figure. **b) Number of inferred contigs with enriched pathways in LECA** (first column). The same graph was randomized 100 times with gene swapping and the number of contigs with a detected GO enrichment is displayed for each randomized graph (second column). **c) Proportion of the adjacencies inferred in LECA conserved in extant eukaryotic genomes**. Each dot represents an extant eukaryotic genome from the OMA database. The x-axis tells whether the genome corresponds to an Animal (Metazoa), a Fungi (Fungi), a Plant (Viridiplantae or Rhodophyta) or a Protist (all other Eukaryotic clades). The y axis gives the proportion of LECA’s adjacencies conserved in the genome. Each genome is colored according to its estimated gene content completeness score, as assessed by OMArk. Each boxplot gives the median, the first and third quartiles (the two hinges). The upper whisker extends from the hinge to the largest value no further than 1.5 * IQR from the hinge (where IQR is the inter-quartile range, or distance between the first and third quartiles) and the lower whisker extends from the hinge to the smallest value at most 1.5 * IQR of the hinge.

### Dating gene adjacencies with EdgeHOG

One novel feature of EdgeHOG is the ability to assess the age of gene adjacencies of extant and ancestral genomes, *i.e*. indicating the last common ancestor in which each adjacency is inferred to have emerged. With it, it is possible to identify patterns of conservation–or lack of– in chromosomal organization over time. We inferred the last common ancestor (clade of origin) of all adjacencies for all eukaryotic genomes in our dataset. In order to be able to compare these data across clades, we dated the adjacencies based on the estimated age of the common ancestor of the corresponding using the TimeTree^26^ resource (**Figure 6** and **Supplementary Data 1**). We observed intriguing patterns of chromosomal evolution.

**Figure 6.**
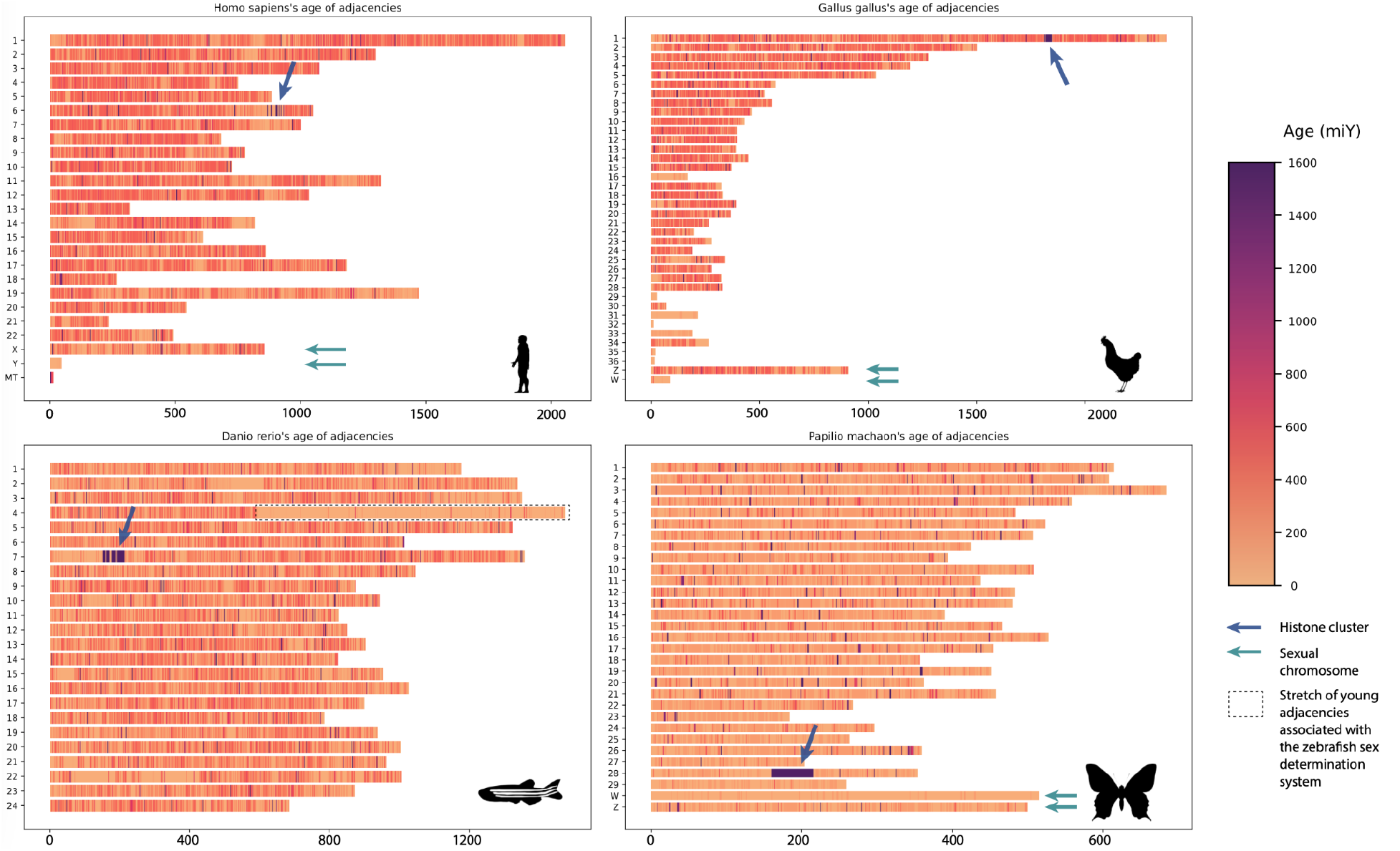
Estimated age of gene adjacencies within chromosomes of *Homo sapiens, Gallus gallus, Danio rerio* and *Papilio machaon*. Each subfigure corresponds to the karyotype of a given species. The y-axis gives the name of the chromosome while the x-axis the number of gene adjacencies. Each chromosome is represented as a stack of gene adjacencies, each colored according to its estimated age in millions of years. The dark blue arrows indicate blocks of adjacencies dated to the eukaryotic common ancestor and enriched in adjacencies between histone genes. The light teal arrows indicate sexual chromosomes, essentially composed of very recent adjacencies, particularly the heteromorphic chromosome (e.g. Y human chromosome).

First, in metazoan genomes, we identified blocks of adjacencies dating from around 1.5 billion years and estimated to have existed in the eukaryotic common ancestor (**Figure 6**, blue arrows). Those clusters of eukaryotic adjacencies involved mainly genes coding for one of the four subunits of the histone octamers (H2A, H2B, H3, H4) and histone linkers (H1/H5). Adjacencies between different histone subunits (hereafter referred to as ‘histone adjacencies’) are observed in species of most of the major eukaryotic clade, which is evidence that these genes existed next to one another in the eukaryotic common ancestor. However, the proportion of those histone adjacencies among all adjacencies was higher in extant Metazoa than in other clades (**Figure S2a**). This may be in part explained by metazoans having more copies of histone genes than most other eukaryotes, although this relationship is not observed in plants despite them having many histone copies as well (**Figure S2b**). Furthermore, the organization of these genes in ‘histone clusters’—occurrences of more than 4 of those adjacencies within close proximity—are observed in many metazoan species but in very few other clades (**Figure S2c**). This points to an organization of histone genes specific to metazoan genomes although the number of the size of those clusters varies between species varying from 12 clusters of 14.75 adjacencies on average in *Bufo bufo* and one cluster of 109 adjacencies in *Drosophila melanogaster* (**Figure S2**). These patterns reveal that an old gene organization pattern—the co-localization of clusters subunit in one locus—took a new form in animals.

The other striking pattern we found while investigating age of adjacencies across the genome related to sex chromosomes (Teal arrows in **Figure 6**). Heteromorphic sex chromosomes are pairs of homologous chromosomes that are morphologically distinct from one another, with one of them carrying a sex determination locus. They are traditionally called X and Y in species where males are heterogametic (XY) and females autogametic (XX), and Z and W when the opposite occurs (ZZ males and ZW females). These systems have been independently acquired multiple times^27^.

In our dataset, heteromorphic sex chromosomes stand out as having younger adjacencies than other chromosomes. We analyzed all species with named sex heterochromosomes in our dataset to investigate this pattern. To do this, for each species, we compared adjacency ages in the sexual chromosome against the distribution of adjacency ages in other chromosomes, and we tested that these two distributions are equivalent with a one sided Mann-Whitney U-test, with the alternative being the age of adjacency in the sexual chromosome is lower

This difference is strong and significant in Y chromosomes (215MiY average difference in 35 mammals, 19.3MiY average difference in diptera) and more moderate but significant for X chromosomes for mammals and diptera but not significantly in nematodes (84.4MiY average difference in 35 mammals, 24.3 average difference in 4 diptera). This pattern was also detected for W/Z systems with the W chromosome having significantly younger adjacencies in the 6 birds in our dataset (308.5MiY), the one fish (199.8MiY) and the one butterfly (122.1MiY) (Per species results and P-value reported in **Table S3**). In contrast, the age of adjacencies for Z chromosomes was significantly smaller than in other chromosomes only in the one butterfly (19.3 MiY younger in average) and some but not all birds while not at all in lizards. In all cases the difference in age was smaller than for the Z chromosomes (**Table S3**). We also noted a pattern of younger adjacencies in the right arm of chromosome 4 in Zebrafish, where the sex determination system is located, but failed to detect it in other fish species with homomorphic sex chromosomes. It is thus unclear that this is a feature of sex determination outside of heteromorphic sex chromosomes. Nevertheless, our observations are evidence that those chromosomes, in particular the one copy existing in minority, undergo different evolutionary trajectories than autosomes in terms of gene organization at least in metazoa. Our automatic dating of adjacencies allowed us to detect this pattern across a wide taxonomic range.

## Discussion

EdgeHOG unlocks key applications in comparative genomics. For instance, it enables tracking genomic rearrangements along a species phylogeny, identifying conserved gene clusters in clades of interest, and improving the assembly of extant genomes by integrating gene order knowledge from other species. Additionally, knowledge of ancestral gene orders can enhance orthology inference by identifying highly divergent orthologs using neighbor genes. The ability to infer gene order conservation across the tree of life also facilitates the comparative genomics of fast-evolving intergenic regions, potentially identifying orthologous regulatory elements using syntenic genes to bracket non-coding regions.

One area of particular interest lies in studying the functional evolution of gene clusters and pathways, offering insights into the conservation and divergence of biological functions across different species. As illustrated in our analyses of LECA’s ancestral contigs, edgeHOG can identify conserved functional gene clusters, such as those involved in essential biological processes like metabolism, DNA repair, and stress responses. This information reveals how these functions have been maintained or adapted throughout evolution. Additionally, the identification of conserved gene neighborhoods can highlight potential new targets for functional studies, as genes located within the same neighborhood can be co-regulated or functionally related.

To perform large-scale inferences, edgeHOG assumes that an adjacency between homologous genes across genomes has been inherited from their last common ancestor. However, such shared adjacencies may result from horizontal gene transfer or independent genomic rearrangements. This assumption can cause erroneous inferences, such as inferring contigs of photosynthetic genes in the ancestral eukaryotic genome due to the cyanobacterial ancestry of chloroplasts in plants, Rhodophyta, or SAR. Therefore, results from edgeHOG in clades with reticulated evolutionary histories need cautious interpretation. Mitigation strategies include using high-confidence HOGs with high completeness scores or removing dubious HOGs that imply numerous gene loss events. EdgeHOG also provides a confidence weight for each ancestral adjacency, reflecting the proportion of extant species supporting it.

The linearized ancestral genomes reconstructed by edgeHOG are optimized for microsynteny, making them less suitable for reconstructing macrosynteny, such as ancestral karyotypes. While they can propose reconstructions of contigs of hundreds to dozens of genes in ancestral species, a single missing adjacency can prevent identifying two neighboring contigs as part of the same chromosome. Integrating microsynteny contigs into a context of ancestral karyotypes would be a logical extension for future versions of edgeHOG. For now, tools like DESCHRAMBLER^6^, which are optimized to maximize the contiguity between ancestral genes, may be more effective at this task.

Our benchmarks consistently show that edgeHOG meets and even slightly exceeds the high standards established by AGORA in recall and precision, with more favorable scaling behavior. This represents an important milestone in comparative genomics. As a key component of the OMA browser’s ecosystem, we anticipate several improvements in future versions, such as predicting the orientation of ancestral genes or modeling lateral gene transfer.

Combining the temporal dimension of gene repertoire evolution with the spatial dimension of gene order evolution provides a comprehensive understanding of genome organization and evolutionary dynamics. This integration opens up more varied applications and advances our knowledge of genome evolution.

## Methods

### Algorithm

#### Extant synteny graphs

PyHam^19^ is called with the orthoXML file of HOGs and the guide species tree (in newick) to model the lineage of all extant genes of all input genomes (**Figure 1**). A lineage is a graph which connects a gene to its parental gene in the upper internal level of the species tree, up until the level of the rootHOG, namely the last common ancestor (LCA) in which the gene is inferred to have emerged. Extant genes not assigned to a HOG are singletons. The genomic coordinates of protein-coding genes in extant genomes are then extracted from either input GFF files or from the HDF5 file of the OMA browser to initialize a linear synteny graph for each leaf of the phylogeny. In such a graph, a node is an extant gene, an edge connects two adjacent genes and each connected component is a chromosome/plasmid/contig (**Figure 2a**).

#### Ancestral synteny graphs

The bottom-up phase propagates edges of synteny graphs from the leaves to the root of the species tree, creating a synteny network at each level of the species tree. An ancestral synteny network is first initialized as a collection of ancestral genes inferred at the current level of the phylogeny. A copy of the synteny graph of each child is then transformed by removing any gene of degree 2 with no parent and reconnecting its two neighbors with a weight equal to the minimal weight between the removed gene and its previous two neighbors (in case the reconnected edge was already present in the network, this minimal weight is added to the initial edge weight). This operation accounts for the insertion of a gained gene between two ancestral genes. By default, only two consecutive reconnections are allowed, but this can be changed by the user, *e.g*. if one expects events like integrations of a mobile genetic element into the chromosome. Edges connecting genes with a distinct parental gene in each transformed child graph are then propagated to the ancestral synteny network (in case the edge was already propagated from another child, its weight is incremented with the weight of the edge in the current child) (**Figure 2a**).

#### Parsimonious ancestral synteny graphs

The top-down phase then prunes edges from synteny networks that are not justified by parsimony, from the root to the leaves of the species tree. At any visited level, the algorithm flags edges whose distribution in children levels and/or in the parental level are supportive that the visited level is either the LCA or a descendant of the LCA in which the edge is thought to have emerged. These correspond to edges present in the propagated network of at least two children, or present in the propagated network of one child and whose propagated form is present in the previously pruned network of the parent. In case two adjacent paralogs result from a duplication at the current level, the propagated form of the edge essentially is the single copy parental gene predating the duplication event. Edges not fulfilling the above conditions are removed from the current synteny network (**Figure 2b**).

#### Linearized graphs

The linearization phase removes edges from each synteny network until the degree of all genes is <= 2. First, conflicting genes (degree > 2) are ordered such that easy-to-resolve genes are treated before the hardest cases. This is done by sorting conflicting nodes by decreasing order of node weight (criterion 2), then by increasing order of degree (criterion 1) (**Figure 2c**). Neighbors of conflicting nodes are then sorted by increasing order of cumulative edge weights in the linear path passing by the visited neighbor and stopping when a node of degree != 2 is met (criterion 2), then by increasing order of the weight of their edge to the conflicting node (criterion 1). This yields a priority list of edges to remove until the degree of the conflicting node becomes 2 (**Figure 2c**). Accounting for the weight of the linear path passing by each neighbor ensures that in cases where multiple neighbors are connected to a conflicting node with the same weight, the one estimated to maximize the weight of the connected component will be prioritized. Yet, because the notion of best neighbor between two genes may not be reciprocal, this may lead to removal of edges between genes that will end up being singletons or terminal nodes of linear contigs at the end of the linearization process. Therefore, a priority list of removed edges to reintroduce is finally created based on the weight that the linear connected component would have if the removed edge was reintroduced in the graph (criterion 2), and based on the weight of the edge itself (criterion 1). Of note, an edge can be reintroduced in the graph only if the two genes still have a degree <2 after the reintroduction of higher-priority edges from the list.

### Benchmarking: preparation of input data

The full dataset of input data for benchmarking is available in **Supplementary Data 2**. The 100 simulated lineages datasets were generated with ALF ^20^, with a low mutational rate (mutRate := 30) to facilitate the downstream detection of orthologs and thus minimize biases inherent to orthology misinferences in the benchmarking of ancestral gene adjacencies prediction. The full list of parameters regarding the compositions of genomes and rates of gene duplications, gene losses and genomic rearrangements are available in **Supplementary Data 2**. The output of ALF is a species tree with a set of ancestral and extant genomes of more or less 400 protein-coding genes (depending on gene losses and duplications) corresponding to all nodes of the species tree. The YGOB dataset (v7-Aug2012) was downloaded from the following url: http://ygob.ucd.ie/. For the OMA Vertebrata dataset, the species tree was obtained by pruning the tree of the Vertebrata clade in OMA using the set of 50 chosen genomes. HOGs were derived from the tree and the all-vs-all of the 50 chosen genomes, exported directly from the OMA browser. For each aforementioned dataset, the preprocessing procedure was the same. First, the OMA Standalone software was called to infer the HOGs from the guide species tree and the proteomes of extant species. HOGs were then converted to a forest of reconciled gene trees, the input format for AGORA. GFF files (for edgeHOG) and lists of ordered genes (for AGORA) were generated from the known order of protein-coding genes in extant genomes for the ALF datasets and the YGOB dataset. For the Vertebrata datasets, the gene order of extant genomes was directly loaded from the HDF5 file of OMA (for the 10 masked species, each gene was considered as a singleton) and converted into lists for AGORA.

### Benchmarking

EdgeHOG and AGORA (default workflow) were called with default parameters against the aforementioned input datasets. For the genome simulation benchmark, inferred adjacencies at each internal level of the species tree were compared to true adjacencies in the corresponding known ancestral genome output by ALF. For the YGOB benchmark, inferred adjacencies at the root of the species tree were compared to YGOB-curated adjacencies in this ancestor. For the masked Vertebrata species benchmarks, any adjacency between two genes in the direct ancestor of a masked extant species were propagated in the masked genome only if the two ancestral genes had each a unique descending gene in the masked genome (no descending paralogs). Projected adjacencies in masked extant genomes were then compared to real unmasked adjacencies. For the simulated lineages and YGOB benchmarks, comparing ancestral adjacencies required to perform a mapping of a modeled ancestral gene (HOG_id in edgeHOG, family_id in Agora) to the corresponding “real” ancestral gene disclosed by ALF and YGOB. This mapping was done based on the maximal number of descending extant genes in common. For each benchmark, the recall score was computed as 100 * TP / (TP + FN) and the precision as 100 * TP / (TP + FP) where TP is the number of true positive adjacencies (correctly inferred adjacencies), FN is the number of false negative adjacencies (missed adjacencies) and FP is the number of false positive adjacencies (misinferred adjacencies).

### Functional analysis of ancestral contigs at the Eukaryota level

The ancestral genome of LECA has been inferred from the Nov2022 release of the OMA database. For each HOG (ancestral gene) defined at the Eukaryota level, the ancestral GO terms were defined as the union of the GO terms of its descending extant genes. The Gene Ontology Enrichment Analysis (GOEA) of each contig was performed with goatools ^28^, using the set of HOGs composing the contig as foreground and all HOGs defined at the Eukaryota level as background. Enriched GO terms were defined as those yielding a Bonferroni p-value adjusted for multiple testing < 0.05 (Fisher’s exact test). Randomized graphs were generated by swapping HOGs among the collection of contigs, which affected only the gene content of contigs and not their topology. The landscape of contigs were annotated and visualized with Cytoscape ^29^.

### Proportion of LECA’s adjacencies conserved in extant eukaryotes

Using pyHAM, we obtained the list of all descendant genes in each species for each HOG present on a contig inferred at the Eukaryota level in the current release of the OMA database. Then for each ancestral adjacency in LECA, we visited the extant synteny graph of each species in which the two LECA’s ancestral genes both have one or more descendant genes. If an extant adjacency between descendant genes of the two LECA’s genes was found, we considered that the extant adjacency was conserved in the assessed species.

### Dating genes of adjacencies

The taxon of origin of adjacencies between genes in extant and ancestral genes were obtained using the date_edges option in EdgeHOG. The estimated age of each taxon was obtained using TimeTree^26^. We first obtained a TimeTree for the species in the OMA Database using the “Load a List of Species” on https://timetree.org. As the TImeTree database does not contain all species present in the OMA Database, we first considered a reduced OMA Taxonomy containing only species shared with TimeTree. We then attributed an age of all non-conflicting internal nodes between OMA and TImeTree corresponding to the distance between any leaf and this internal node. Finally, we attributed an age to 0 to any leaf in the OMA Taxonomy. For any node left with no age at this point, we assigned an age corresponding to the average of the age of its ancestor and the age of the oldest of its children as a default.

#### Histone adjacencies cluster

We noted as representative of histone gene members of the HOGs. We defined histone adjacency clusters as any group of genes in which there were more than 4 adjacencies between histone genes of distinct HOGs, and in which there were less than 10 genes between any of those adjacencies. Cluster “size” was defined as the number of histone adjacencies found within the cluster.

#### Sex chromosomes

In order to compare sex chromosomes specificity across our dataset, we selected the ones for which this information was available in a non-ambiguous way through the OMA Database. To do this, we selected all genomes in our data for which chromosomes were identified by a number, or a letter within the X, Y, Z, W system. From this selection, we removed Fungi genomes for which represented a roman numeral and not a sex chromosome. From the selected genomes, we considered only canonically named chromosomes (numeric identifier or letters) to discount incomplete contigs and scaffolds. Finally, all comparisons were done between one of the sex chromosomes against all the other complete chromosomes. We used one-sided Mann-Whitney tests to test whether the distribution of adjacency ages was similar between the sex and the other chromosomes, with the alternative hypotheses being the sex chromosome having younger adjacencies than the others.

## Code availability

EdgeHOG is free open-source software (MIT License) available at https://github.com/DessimozLab/edgeHOG. Code and scripts used in the analyses of this paper are available in **Supplementary Data 2**.

## Supplementary materials

- **Supplementary Figure 1**: Selection of the 10 masked genomes and the 40 other representative genomes within the Vertebrata clade in OMA + Benchmarking of EdgeHOG with the full Vertebrata clade composed of 156 species
- **Supplementary Figure 2**: Histone clusters in eukaryotic clades
- **Supplementary Table 1**: Functional annotation of ancestral contigs in LECA
- **Supplementary Table 2**: Conservation of LECA’s adjacencies in extant eukaryotes
- **Supplementary Table 3**: Average age of adjacencies in sex chromosomes vs other chromosomes
- **Supplementary Data 1**: Plots with dated adjacencies for 706 eukaryotic genomes: https://sibcloud-my.sharepoint.com/:u:/g/personal/christophe_dessimoz_sib_swiss/ESSJe-O1fXJCiBohKN4-4J4BT2y0HPHdEkWZRz0K_kl2UQ?e=DS83Y0 (*temporary location, this will eventually be released as a data archive, likely on Figshare*)
- **Supplementary Data 2** (https://figshare.com/s/a0b482b1d6e236c43005): This repository contains all the scripts and datasets (simulations, YGOB, OMA Vertebrata species) used for the benchmarking of EdgeHOG. It all also contains the data and scripts used for downstream analyses, i.e the functional annotation of reconstructed contigs in LECA, the study of the conservation of LECA’s adjacencies in extant eukaryotes and the dating of gene adjacencies in extant eukaryotes

## Author contribution

CB designed the top-down and linearization phases of edgeHOG, implemented the software, performed the benchmarking, conducted the downstream analyses and wrote the manuscript with input from all coauthors. NBRK contributed to the design and the code of the ALF and YGOB benchmarks. KJG designed the bottom-up phase of edgeHOG. AWV contributed to the ancestral GO enrichment analysis. CT implemented pyHAM and wrote the code to explore and visualize ancestral gene orders on the OMA browser. YN assessed the quality of ancestral adjacency reconstructions, and designed, performed, and interpreted the adjacency dating analyses. NG participated in the design of the full study, contributed to the manuscript, and contributed to project supervision. AA contributed to the code of the preprocessing, bottom-up and outputting steps of edgeHOG, contributed to the design and implementation of the benchmarking protocoles, and in the integration of edgeHOG in the ecosystem of tools of the OMA browser. CD conceptualized and supervised the project.

## Inclusion & Ethics

Our study relies on publicly available datasets and our results are broadly accessible and reproducible. We analyzed a diverse array of genomes from bacteria, archaea, and eukaryotes, encompassing a wide range of species and genetic diversity. This comprehensive approach ensures our findings are relevant across various biological contexts.

Ethical considerations guided our research. All computational analyses were conducted using publicly available genomic data from the OMA orthology database and other established sources, which can be traced back to the original data sources (https://omabrowser.org/All/oma-species.txt). No new genomic data was sourced from private or sensitive origins. We have committed to open science by making edgeHOG available as open-source software on PyPI and GitHub, ensuring free access and further development by the scientific community. Additionally, scripts and datasets used in our analyses are available on Figshare to promote reproducibility.

