## Supplementary Figures for "EdgeHOG: fine-grained ancestral gene order inference at tree-of-life scale"

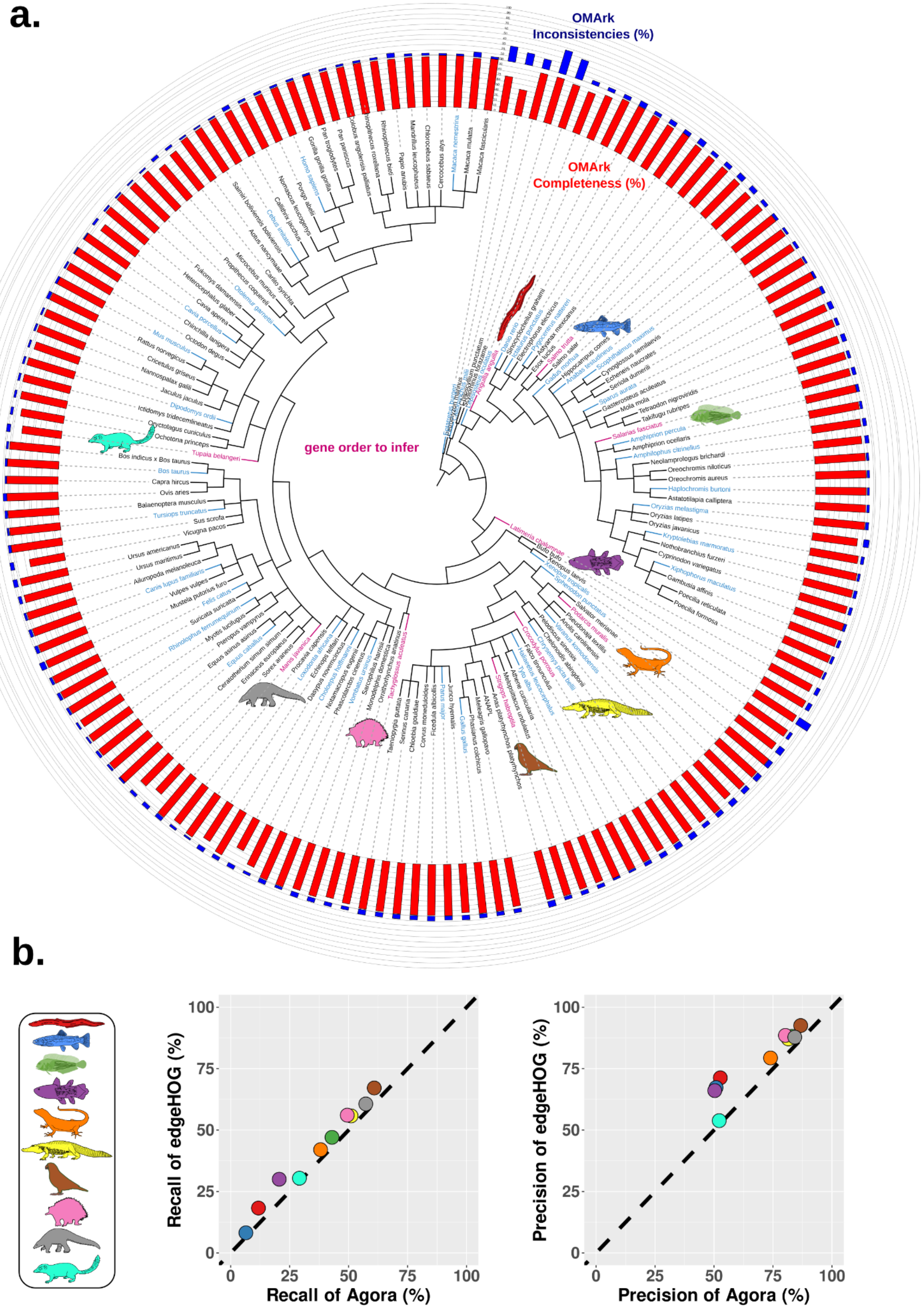
a.
OMArk Inconsistencies (%)
OMArk Completeness (%)
gene order to infer
b.
100
75
50
25
0
Recall of edgeHOG (%)
Recall of Agora (%)
Precision of edgeHOG (%)
Precision of Agora (%)

**Figure S1. a) Selection of the 10 masked genomes** (pink) **and the 40 other representative genomes within the Vertebrata clade in OMA** (blue)**.** The bars give the OMArk-assessed completeness and the consistency of gene repertoire of each genome relative to the closest species in OMA. **b) Benchmarking with the full Vertebrata clade composed of 156 species**

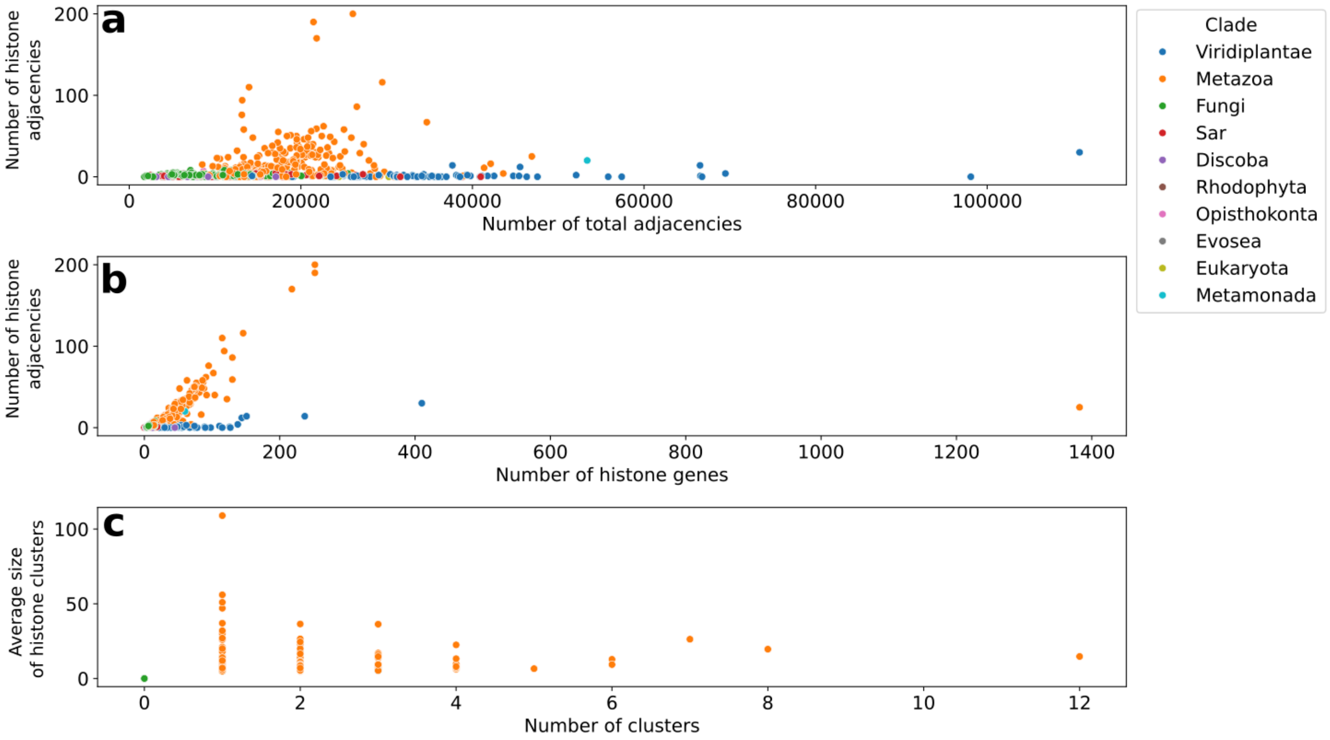


`

**Figure S2**. Histone clusters in eukaryotic clades. **a)** Adjacencies between histone genes are more prevalent in metazoa. Number of adjacencies between distinct histone genes as a function of number of total genewise adjacencies. **b)** Number of histone adjacencies is proportional to number of histone genes in metazoa. Number of adjacencies between distinct histone genes as a function of the number of histone genes. **c)** Organization of histones in gene clusters in metazoa. Number of distinct histone clusters (x) and average number of adjacencies between histone genes within it (y). Colors indicate the eukaryotic clade to which each species belongs to.
